## supplementary_section for "Stochastic modeling of a gene regulatory network driving B cell development in germinal centers"

### A Supplementary Material

#### A.1 Mathematical reduction of the stochastic model and estimation of the initial activation rate

From Martinez et al. [3], one can see that IRF4 is a crucial node and its autoactivation reaction is responsible for the bistability switch from the GC to PB_PC during B cell differentiation. Notably, equation (3) which describes the dynamics of IRF4, only depends on IRF4. Based on these observations, we have decided to use IRF4 as a connecting edge between models (4)-(6) and (1)-(3).
Starting from the stochastic model (4)-(6) written for IRF4, we reduced it to an ODE model by making a simplifying assumption. We substitute the stochastic process $E(t)$ by its mean value $\langle E(t)\rangle$, so

$$\begin{cases} \dfrac{dM_{\text{IRF4}}}{dt} &= s_{0,\,\text{IRF4}}\langle E_{\text{IRF4}}(t)\rangle - d_{0,\,\text{IRF4}}M_{\text{IRF4}}(t), \\ \dfrac{dP_{\text{IRF4}}}{dt} &= s_{1,\,\text{IRF4}}M_{\text{IRF4}}(t) - d_{1,\,\text{IRF4}}P_{\text{IRF4}}(t). \end{cases} \tag{SF.1}$$

We are looking for parameter values of system (SF.1) that will allow to reproduce the same behavior than the kinetic model (1)-(3) (i.e. two steady states). At steady state, system (SF.1) with $\frac{dM}{dt} = 0$ and $\frac{dP}{dt} = 0$ leads to:

$$P_{\text{IRF4}}(t) = \frac{s_{1,\,\text{IRF4}}s_{0,\,\text{IRF4}}\langle E_{\text{IRF4}}(t)\rangle}{d_{0,\,\text{IRF4}}d_{1,\,\text{IRF4}}}. \tag{SF.2}$$

Introducing the new variable

$$c = \frac{s_{1,\,\text{IRF4}}s_{0,\,\text{IRF4}}}{d_{1,\,\text{IRF4}}d_{0,\,\text{IRF4}}}$$

we can write (SF.2) as

$$P_{\text{IRF4}}(t) = c\langle E_{\text{IRF4}}(t)\rangle. \tag{SF.3}$$

In Martinez et al. [3], IRF4 behavior was described by equation (3), that is

$$\frac{dr}{dt} = \mu_r + \sigma_r\frac{r^2}{k_r^2 + r^2} + CD40 - \lambda_r r,$$

i.e. with our notation it can be written as:

$$\frac{dp_{\text{IRF4}}}{dt} = \mu_{\text{IRF4}} + \sigma_{\text{IRF4}}\frac{p_{\text{IRF4}}^2}{k_{\text{IRF4}}^2 + p_{\text{IRF4}}^2} + CD40 - \lambda_{\text{IRF4}}\,p_{\text{IRF4}}. \tag{SF.5}$$

Assuming that CD40 $= 0$ at the beginning of the simulation, that equation (SF.5) is at the steady state, that $\lambda_{\text{IRF4}}$ is a degradation rate of protein for IRF4 and using (SF.3), we write (SF.5) as:

$$\mu_{\text{IRF4}} + \sigma_{\text{IRF4}}\frac{c^2(\langle E_{\text{IRF4}}(t)\rangle)^2}{k_{\text{IRF4}}^2 + c^2(\langle E_{\text{IRF4}}(t)\rangle)^2} - cd_{1,\,\text{IRF4}}\langle E_{\text{IRF4}}(t)\rangle = 0. \tag{SF.6}$$

Solving equation (SF.6) in terms of $\langle E_{\text{IRF4}}(t)\rangle$ leads to

$$0 = c^3 d_{1,\,\text{IRF4}}(\langle E_{\text{IRF4}}(t)\rangle)^3 - (\mu_{\text{IRF4}} + \sigma_{\text{IRF4}})c^2(\langle E_{\text{IRF4}}(t)\rangle)^2 \\ + c d_{1,\,\text{IRF4}} k^2_{\text{IRF4}}(\langle E_{\text{IRF4}}(t)\rangle) - \mu_{\text{IRF4}} k^2_{\text{IRF4}}$$

and can be simplified in the form:

$$a'(\langle E_{\text{IRF4}}(t)\rangle)^3 - b'(\langle E_{\text{IRF4}}(t)\rangle)^2 + c'(\langle E_{\text{IRF4}}(t)\rangle) + d' = 0 \tag{SF.8}$$

where

$$\begin{cases} a' = c^3 d_{1,\,\text{IRF4}} \\ b' = (\mu_{\text{IRF4}} + \sigma_{\text{IRF4}})c^2 \\ c' = c d_{1,\,\text{IRF4}} k^2_{\text{IRF4}} \\ d' = -\mu_{\text{IRF4}} k^2_{\text{IRF4}} \end{cases}$$

Because parameters $a', b', c'$ are positive and $d'$ is negative, there is at least one positive root to (SF.8). Further, we fitted the parameters $\mu_{\text{IRF4}}, \sigma_{\text{IRF4}}, k_{\text{IRF4}}$, applying fitting procedure from Martinez et al. [3] and using the experimental data accession no. GSE 12195 (see Tables 2-5), and we found that $E_{\text{IRF4}}(t_{\text{init}})$ value which would correspond to a bistable regime of system (1)-(3) is:

$$E_{\text{IRF4}}(t_{\text{init}}) = 1.7 \times 10^{-3}.$$

We also know that

$$\langle E_{\text{IRF4}}\rangle = \frac{k_{\text{on}}}{k_{\text{on}} + k_{\text{off}}},$$

so assuming that at initial time $t = t_{\text{init}}$, $k_{\text{on}} << k_{\text{off}}$ and $k_{\text{on}} = \alpha k_{\text{off}}$, with $\alpha << 1$, one can define $\alpha = E_{\text{IRF4}}/(1 - E_{\text{IRF4}}) = 1.7 \times 10^{-3}$. Assuming $k_{\text{off}} \approx 1$ ($k_{off,\,\text{init},\,\text{IRF4}} \approx 1$) allows to estimate the value of $k_{on}$ for IRF4, which should keep reduced model (11) in a two steady state regime:

$$k_{on,\,\text{IRF4}} = 1.7 \times 10^{-3} \tag{SF.9}$$

Further, we called the value (SF.9), the initial value $k_{on,\,\text{init}}$ for IRF4.

#### A.2 Supplementary Figures

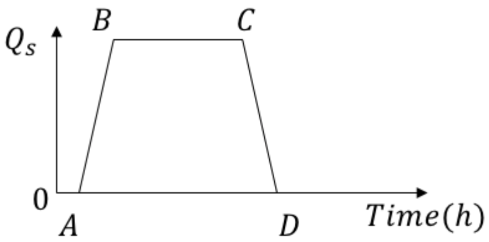


**Supplementary Figure S1.** The scheme of application of the stimuli $Q_s$, where $s \in \{BCR, CD40\}$. Stimuli $Q_s$ were implemented in three steps: $AB$ - linear increase ($t_{\text{BCR}} \in [0.5h; 1.5h]; t_{\text{CD40}} \in [35h; 36h]$), $BC$ - stable stimulus ($t_{\text{BCR}} \in [1.5h; 24h]$; $t_{\text{CD40}} \in [36h; 60h]$), $CD$ - linear decrease ($t_{\text{BCR}} \in [24h; 25h]; t_{\text{CD40}} \in [60h; 61h]$).

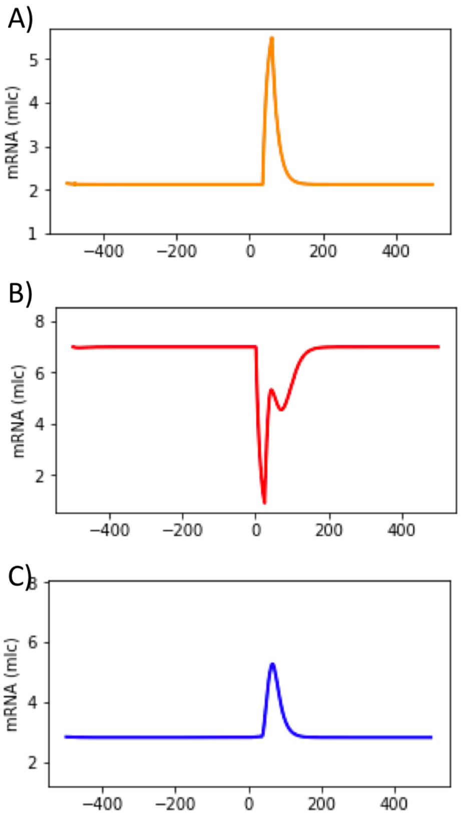


**Supplementary Figure S2.** Absence of bistability in model (11), due to random choice of parameter values. The average behavior of solutions of model (11) are displayed: IRF4 (A), BCL6 (B) and BLIMP1 (C) proteins levels, as depicted in Figure 1A). BCR stimulus was applied from 0h until 25h and CD40 stimulus from 35h until 60h. Parameters used for (11) are listed in Supplementary Table S1 and $k_{on,\,\text{init},\,\text{IRF4}} = 0.1$.

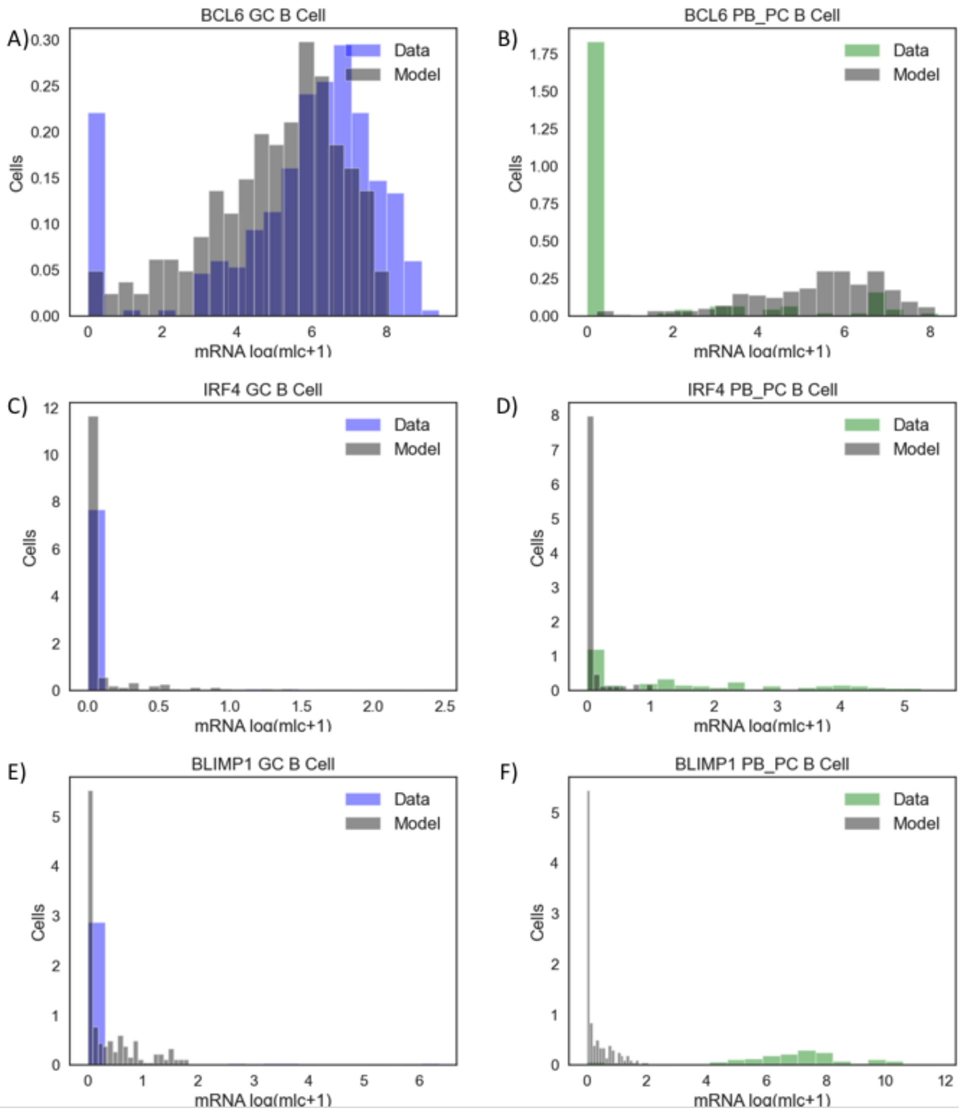


**Supplementary Figure S3.** The histograms of model-generated and experimental mRNA counts of BCL6, IRF4, BLIMP1 at GC and PB_PC stages. The subgraphs A, C, E represent log (molecule+1) transformed SC with BCL6, IRF4 and BLIMP1 compared between the model estimations at GC stage (grey) *vs* the experimental data from GC B cells (blue). The subgraphs B, D, F represent log (molecule+1) transformed SC with BCL6, IRF4 and BLIMP1 compared between the model estimations at PB_PC stage (grey) *vs* the experimental data from PB_PC cells (green). Simulation of 200 SC was used based on the parameter set selected after automatized parameter screening strategy (see Tables 2-5, version II).

#### A.3 Supplementary Tables

| Parameter | Values | Parameter | Values |
|---|---|---|---|
| $H_{11}$ | 1 | $s_{0,\mathrm{BCL6}}$ | 6.5 |
| $H_{21}$ | 0.1 | $s_{0,\mathrm{IRF4}}$ | 2 |
| $H_{31}$ | 1 | $s_{0,\mathrm{BLIMP1}}$ | 6.5 |
| $H_{12}$ | 1 | $d_{0,\mathrm{BCL6}}$ | 0.05 |
| $H_{22}$ | 0.01 | $d_{0,\mathrm{IRF4}}$ | 0.05 |
| $H_{32}$ | 1 | $d_{0,\mathrm{BLIMP1}}$ | 0.1733 |
| $H_{13}$ | 0.1 | $s_{1,\mathrm{BCL6}}$ | 100 |
| $H_{23}$ | 0.001 | $s_{1,\mathrm{IRF4}}$ | 160 |
| $H_{33}$ | 1 | $s_{1,\mathrm{BLIMP1}}$ | 40 |
| $H_{\mathrm{BCR},1}$ | 0.01 | $d_{1,\mathrm{BCL6}}$ | 0.138 |
| $H_{\mathrm{CD40},2}$ | 1 | $d_{1,\mathrm{IRF4}}$ | 0.173 |
| $\theta_{11}$ | -0.2 | $d_{1,\mathrm{BLIMP1}}$ | 0.173 |
| $\theta_{21}$ | -10 | $k_{\mathrm{on,\,init,\,BCL6}}$ | 0.1 |
| $\theta_{31}$ | -2 | $k_{\mathrm{on,\,init,\,IRF4}}$ | 0.1 |
| $\theta_{12}$ | 0 | $k_{\mathrm{on,\,init,\,BLIMP1}}$ | 0.1 |
| $\theta_{22}$ | 8 | $k_{\mathrm{off,\,init,\,BCL6}}$ | 1 |
| $\theta_{32}$ | 0 | $k_{\mathrm{off,\,init,\,IRF4}}$ | 1 |
| $\theta_{13}$ | -1 | $k_{\mathrm{off,\,init,\,BLIMP1}}$ | 1 |
| $\theta_{23}$ | 40 | | |
| $\theta_{33}$ | 0 | | |
| $\theta_{\mathrm{BCR},1}$ | -200 | | |
| $\theta_{\mathrm{CD40},2}$ | 10 | | |

**Supplementary Table S1.** Parameters of System (11) with values accordingly to Bonnaffoux et al. [32].
